# A 3D Clinical Face Phenotype Space of Genetic Syndromes using a Triplet-Based Singular Geometric Autoencoder

**DOI:** 10.1101/2022.12.27.521999

**Authors:** Soha S. Mahdi, Eduarda Caldeira, Harold Matthews, Michiel Vanneste, Nele Nauwelaers, Meng Yuan, Shunwang Gong, Giorgos Bouritsas, Gareth S Baynam, Peter Hammond, Richard Spritz, Ophir D Klein, Michael Bronstein, Benedikt Hallgrimsson, Hilde Peeters, Peter Claes

## Abstract

Clinical diagnosis of syndromes benefits strongly from objective facial phenotyping. This study introduces a novel approach to enhance clinical diagnosis through the development and exploration of a low-dimensional metric space referred to as the clinical face phenotypic space (CFPS). As a facial matching tool for clinical genetics, such CFPS can enhance clinical diagnosis. It helps to interpret facial dysmorphisms of a subject by placing them within the space of known dysmorphisms. In this paper, a triplet loss-based autoencoder developed by geometric deep learning (GDL) is trained using multi-task learning, which combines supervised and unsupervised learning approaches. Experiments are designed to illustrate the following properties of CFPSs that can aid clinicians in narrowing down their search space: a CFPS can 1) classify syndromes accurately, 2) generalize to novel syndromes, and 3) preserve the relatedness of genetic diseases, meaning that clusters of phenotypically similar disorders reflect functional relationships between genes. The proposed model consists of three main components: an encoder based on GDL optimizing distances between groups of individuals in the CFPS, a decoder enhancing classification by reconstructing faces, and a singular value decomposition layer maintaining orthogonality and optimal variance distribution across dimensions. This allows for the selection of an optimal number of CFPS dimensions as well as improving the classification capacity of the CFPS.

## I. Introduction

Human genetic syndromes are rare and diverse and thus, diagnosing them can be a challenging and complex process [1]. Changes in the anatomical structure of the craniofacial region are common in patients with genetic diseases, affecting between 30 and 40% of the patients with this type of condition [2], meaning that a preliminary diagnosis can be performed by analyzing the patient’s facial phenotypical clues. However, this task presents complexities and challenges. On the one side, the spectrum of possible conditions is wide. While some have highly distinguishable facial traits, others share identical or have only subtle dysmorphic clues, which can lead to an incorrect diagnosis. In each syndrome, individuals may exhibit varying levels of severity, leading to differences in their observable characteristics [3]. On the other side, such a broad spectrum of possible conditions and phenotypic variability results in subjective diagnoses, which are highly dependent on the clinician’s level of expertise [4].

To combat these issues, deep learning (DL) approaches have been proposed for the syndrome classification task based on facial phenotypical clues. DL models are known for their ability to learn complex tasks. In the case of genetic syndrome classification, they also provide a more objective way of performing the diagnosis [4], removing the subjectivity imposed by the clinician’s level of expertise. While it has already been shown that artificial intelligence (AI) systems can outperform clinicians by a large margin in the syndrome classification task [5], these models are not flawless. As such, the final diagnosis can benefit from the combined use of clinicians’ expertise and AI tools that efficiently narrow down the disease spectrum that the clinician has to analyze [5], facilitating their task.

A pivotal advancement in the study of genetic syndromes was the introduction of the Clinical Face Phenotype Space (CFPS) by Ferry et al. [6]. Utilizing metric learning techniques, CFPS models facial dysmorphisms in a lower-dimensional latent space, offering a novel approach to understanding these complex syndromes. The CFPS is characterized by three main properties. Firstly, it quantifies phenotypic similarity, grouping patients’ faces based on diagnostically relevant features. Secondly, its ability to generalize allows it to encapsulate dysmorphic syndromes beyond those used in its initial training, making it a versatile tool for exploring and categorizing both known and novel syndromes [7]. Lastly, it reflects the underlying genetic relationships by grouping phenotypically similar disorders, thereby mirroring the functional relationships among their genes [7]. This space not only facilitates rapid comparative analysis among individuals but also aids in proposing hypothetical clinical and molecular diagnoses, thereby streamlining genome wide NGS analyses and guiding targeted sequencing in clinical diagnostics [8].

However, constructing a CFPS is not without challenges. The overlapping features of different syndromes, as well as the significant phenotypic variation within each syndrome, make it difficult to encode facial shapes accurately [9]–[13]. For instance, hypertelorism is a feature shared by both Apert and Wolf-Hirschhorn syndromes [14], [15]. To address these challenges, our proposed model incorporates a supervised metric learner based on a triplet loss function. This component is designed to optimize the CFPS by learning to discriminate between groups while recognizing within-group similarities.

Yet, focusing solely on discriminatory aspects may overlook general facial topology. To counter this, we integrate unsupervised dimensionality reduction techniques, such as autoencoders, with the metric learning encoder. This combination aims to preserve facial topology within the CFPS, ensuring that it captures both diagnostically relevant features and general facial variations. This dual approach allows the model to effectively represent both specific syndromic features and more general patient-to-patient variations, essential for encoding patients with unseen syndromes.

Moreover, our model’s decoder is capable of reconstructing a face from a sampled embedding in the CFPS. However, without a structured approach during the training of this encoder-decoder, reliable sampling from the latent space remains a challenge. To enhance this capability, we introduce a third component: a singular value decomposition (SVD) layer. This layer, first proposed by Nauwelaers et al. [16], imposes orthogonality and optimizes variance distribution across the dimensions of the latent space, enhancing the model’s ability to handle non-linearity and an easier determination of dimensionality.

## II. Materials and Methods

### A. Data

The 3D facial images utilized in this project were sourced from three primary collections:

1. The FaceBase repository^1^, specifically the project “Developing 3D Craniofacial Morphometry Data and Tools to Transform Dysmorphology, FB00000861”. These images were collected from patient support groups in the USA, Canada, and the UK [17], [18].
2. The Health Department of Western Australia. This dataset comprises images collected between 2009 and 2018, primarily through Genetic Services of Western Australia and additionally at Australian hospitals and patient support groups [19].
3. The legacy 3D dysmorphology dataset of Peter Hammond, hosted at KU Leuven, Belgium. This collection includes patients from support groups across the United States, UK, and Italy, spanning from 2002 to 2013. Diagnoses were reported by families and/or clinical geneticists [3].

From these sources, groups with a minimum of 10 individuals were selected for inclusion in the study. The distribution of the data is approximately 59%, 40%, and less than 1% from the first, second, and third sources, respectively. The dataset comprises 3,496 3D facial images, representing 52 different syndromes and a control group of 100 individuals unrelated to the patients with known genetic syndromes. Detailed demographic characteristics of the dataset are presented in Table 1.

**TABLE I:** Data demographics: Syndrome group name, sample size (N), mean and standard deviation of age (M±SD), the sex ratio (female/(male+female)), and the category.

| Name | Size | Age Range | Sex Ratio | Category | Name | Size | Age Range | Sex Ratio | Category |
| --- | --- | --- | --- | --- | --- | --- | --- | --- | --- |
| Williams | 221 | 17.57 ± 13.9 | 0.46 | A | BBS | 87 | 26.33 ± 14.78 | 0.48 | C |
| 22q11.2 Del | 180 | 10.74 ± 6.03 | 0.49 | A | Neurofibromatosis | 85 | 20.18 ± 18.01 | 0.54 | C |
| Wolf Hirschhorn | 155 | 11.03 ± 9.42 | 0.57 | A | Loeys Dietz | 84 | 25.38 ± 17.15 | 0.57 | C |
| Smith Magenis | 129 | 14.32 ± 9.09 | 0.55 | A | Joubert | 75 | 10.57 ± 8.58 | 0.48 | C |
| Down | 117 | 21.64 ± 11.14 | 0.49 | A | Ectodermal Dysplasia | 71 | 15.09 ± 15.32 | 0.28 | C |
| Prader Willi | 96 | 19.34 ± 13.24 | 0.51 | A | Rett | 70 | 13.32 ± 10.54 | 0.89 | C |
| Fragile X | 77 | 17.65 ± 12.56 | 0.3 | A | Cardiofaciocutaneous | 59 | 12.21 ± 8.55 | 0.53 | C |
| Achondroplasia | 70 | 22.62 ± 18.34 | 0.59 | A | Klinefelter | 57 | 22.91 ± 14.58 | 0 | C |
| Rubinstein Taybi | 63 | 13.54 ± 11.73 | 0.52 | A | Mucopolysaccharidosis | 57 | 21.51 ± 13.51 | 0.47 | C |
| Costello | 58 | 12.39 ± 9.24 | 0.66 | A | Alstrom | 52 | 21.28 ± 9.4 | 0.54 | C |
| Cohen | 33 | 18.27 ± 10.46 | 0.52 | A | Fibrodysplasia Ossificans Progressiva | 50 | 21.81 ± 12.37 | 0.56 | C |
| Pitt Hopkins | 29 | 8.53 ± 5.7 | 0.62 | A | Fabry | 48 | 32.37 ± 16.53 | 0.44 | C |
| Pallister Killian | 23 | 9.59 ± 7.19 | 0.26 | A | Sotos | 45 | 17.92 ± 12.22 | 0.49 | C |
| Crouzon | 22 | 10.22 ± 6.27 | 0.55 | A | Russell Silver | 44 | 10.18 ± 10.32 | 0.34 | C |
| Smith Lemli Opitz | 19 | 11.75 ± 7.05 | 0.32 | A | Cockayne | 41 | 12.15 ± 7.37 | 0.44 | C |
| Apert | 13 | 14.55 ± 10.73 | 0.62 | A | Pseudoachondroplasia | 35 | 28.06 ± 20.53 | 0.51 | C |
| Coffin Lowry | 12 | 13.76 ± 9.16 | 0.08 | A | Osteogenesis Imperfecta | 31 | 16.72 ± 14.61 | 0.68 | C |
| Cornelia de Lange | 183 | 12.1 ± 9.14 | 0.54 | B | 1p36 Del | 29 | 8.82 ± 7.83 | 0.62 | C |
| Noonan | 155 | 14.06 ± 12.51 | 0.45 | B | Trisomy 18 | 27 | 8.79 ± 8.71 | 0.85 | C |
| Angelman | 106 | 9.97 ± 7.57 | 0.47 | B | Beckwith Wiedemann | 26 | 9.68 ± 6.66 | 0.42 | C |
| Stickler | 45 | 22.31 ± 17.45 | 0.62 | B | EED CLP | 20 | 23.24 ± 17.71 | 0.65 | C |
| Treacher Collins | 39 | 18.48 ± 13.5 | 0.49 | B | Vander Woude | 16 | 10.16 ± 4.75 | 0.56 | C |
| Kabuki | 37 | 12.09 ± 6.62 | 0.65 | B | Goltz | 14 | 9.4 ± 5.03 | 0.86 | C |
| Coffin Siris | 16 | 12.08 ± 9.65 | 0.63 | B | Rhizo Chondro Punct | 13 | 7.57 ± 5.53 | 0.69 | C |
| 5p Del Cri du Chat | 96 | 14.62 ± 12.91 | 0.58 | C | Zellweger Syndrome | 11 | 7.33 ± 9.49 | 0.09 | C |
| Marfan | 153 | 26.34 ± 16.85 | 0.58 | C | Controls | 100 | 30.94 ± 11.64 | 0.72 | CONTROL |
| Turner | 102 | 24.25 ± 19.03 | 0.98 | C |  |  |  |  |  |

Based on clinical assessment of the available images, two clinical experts (co-authors HP and MV) categorized the syndromic groups into three categories:

a. Genetic conditions diagnosable by typical facial characteristics, which are genetically homogeneous (caused by a single gene or recurrent chromosomal anomaly).
b. Genetic conditions diagnosable by typical facial characteristics but genetically heterogeneous (multiple genes associated with the clinical condition).
c. Conditions usually not diagnosed based on facial features, where facial features are not typically characteristic of the condition.

For syndromes in categories A and B, facial features generally direct clinicians towards a molecular diagnosis. However, the genetic heterogeneity in category B can introduce variability in the phenotype-genotype correlation. Category C encompasses syndromes typically diagnosed based on clinical symptoms other than facial features. Although a distinctive facial gestalt is not a primary diagnostic criterion for these syndromes, it is not definitively absent.

This study received ethical approval from the ethical review board of KU Leuven and University Hospitals Gasthuisberg, Leuven (Approval Numbers: S56392, S60568).

### B. Preprocessing

For pre-processing, after cleaning the raw image by removing hair and ears, a 3D face template was non-rigidly registered to each face using Meshmonk [20]. Each 3D face shape is therefore described as a manifold triangle mesh *F* = (*V, ℰ*, Φ), where 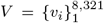 is a 8, 321 *×* 3 dimensional matrix, containing 8,321 3D vertices *v*_*i*_ = (*x*_*i*_, *y*_*i*_, *z*_*i*_) defining the mesh geometry, ℰ and Φ are set of edges and faces which define the mesh topology. ℰ and Φ are fixed since all our meshes have the same topology as the template.

### C. Pipeline Design

This section outlines the design of our computational pipeline for learning a Clinical Face Phenotype Space (CFPS). Our approach begins with a baseline method using Linear Discriminant Analysis (LDA), a well-established supervised linear metric learner and classifier. Beyond this, we propose a more advanced, multi-component metric learner based on geometric deep models, comprising the following elements:

1. A triplet-loss encoder, which is a supervised deep-metric learner. It consists of three identical encoder networks, trained with triplets of data including an anchor (*f*_*a*_), a positive (*f*_*p*_), and a negative sample (*f*_*n*_). In each triplet, the anchor and positive samples belong to the same class, while the anchor and negative samples are from different classes. The network outputs lower-dimensional embeddings for each element of the triplet: (*e*_*a*_, *e*_*p*_, *e*_*n*_) = (*p*_*GE*_ (*f*_*a*_), *p*_*GE*_ (*f*_*p*_), *p*_*GE*_ (*f*_*n*_)).
2. A decoder (*p*_*GD*_) that reconstructs the 3D facial meshes from these embeddings: (*f′*_*a*_, *f′* _*p*_, *f′*_*n*_) = (*p*_*GD*_(*e*_*a*_), *p*_*GD*_(*e*_*p*_), *p*_*GD*_(*e*_*n*_)).
3. An SVD layer to ensure orthogonal dimensions and optimal variance distribution across different CFPS dimensions.

These components are gradually integrated to form our Triplet-based Singular Geometric Autoencoder (TB-SGAE), which utilizes spiral-based geometric architectures. Figure 1 illustrates the main components of this model. In the following sections, we begin with a description of the baseline method. We then methodically detail each component, explaining their individual development and how they integrate to enhance the TB-SGAE, and lastly, we provide the details of our spiral-based geometric autoencoder architecture.

**Fig. 1:**
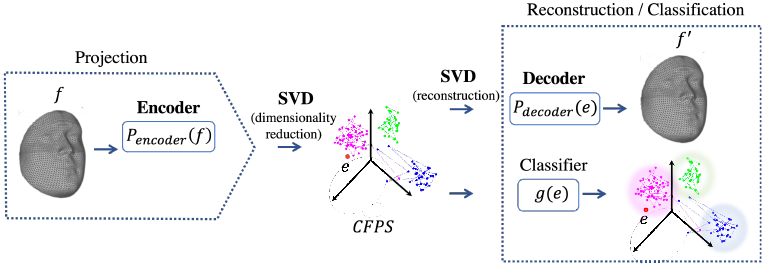
The complete model consists of three main components: a triplet-based encoder, a singular value decomposition (SVD) layer, and a decoder. Projection function *p*_*SGE*_ for geometric model (alternatively *p*_*LDA*_ for baseline) projects a facial mesh f into a facial embedding e in the CFPS. A facial mesh *f*^*′*^ is reconstructed from the embedding e with decoding function *p*_*SGD*_. Note that the reconstruction is not possible within the baseline and TB-GAE. Classification from the embedding space into syndrome groups is performed by a classification function g, which in this work constitutes a balanced K-nearest-neighbor classifier.

#### 1) CFPS based on Linear Discriminant Analysis (LDA)

Linear Discriminant Analysis (LDA) is utilized to maximize the between-class scatter (*S*_*b*_) while minimizing the within-class scatter (*S*_*w*_) for facial data. Due to the high dimensionality of our densely sampled meshes, we applied Principal Component Analysis (PCA) to reduce dimensions and mitigate overfitting in LDA. The first 100 dimensions, preserving 99.16% of data variation, were then used for LDA. We constructed the lower-dimensional space projection using Fisher’s criterion 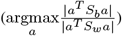, where *a* is the projection matrix. The projected embedding for a facial shape *f* in CFPS is calculated as:

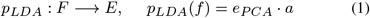

The maximum number of dimensions for LDA was set to the number of classes minus one (52).

#### 2) Triplet-Based Geometric Encoder (TB-GE)

Our Triplet-Based Geometric Encoder (TB-GE) employs spiral convolutional operators for mapping 3D facial meshes to a CFPS. It ensures that the feature representations of patients within the same syndrome group are positioned closer compared to those from different groups. The encoder, denoted as *p*_*GE*_, maps an input mesh *f* ∈ *F* to a low-dimensional embedding *e* ∈ *E* in CFPS:

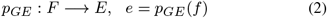

The triplet-loss function for training the TB-GE is defined as:

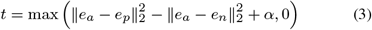

where *α* is the margin between positive and negative samples, set to 0.2 as per Schroff et al. [21].

#### 3) Geometric Decoder

The geometric decoder (GD) function *p*_*GD*_ reconstructs a facial mesh *f*^*′*^ from an embedding *e*:

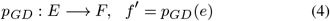

The Triplet-Based Geometric Autoencoder (TB-GAE) is formulated as:

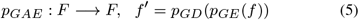

The reconstruction loss for 3D facial shapes is the mean absolute error between vertices of the input shape *f* and the reconstructed output *f′*, averaged over the dataset:

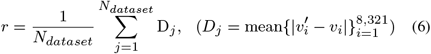

The final loss function for training the TB-GAE combines reconstruction and triplet losses:

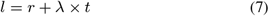

where *λ* balances the two loss components and is set to 0.1 to ensure equal contribution of both losses.

#### 4) SVD Layer: Decorrelation of CFPS Dimensions

To achieve orthogonal dimensions in CFPS, Singular Value Decomposition (SVD) is applied to the embeddings set *E*: *US*Λ^*T*^ = *E*. Here, *S* is a diagonal matrix of singular values, *U* and Λ contain left and right singular vectors, respectively. The projection function *p*_*SGE*_ is defined as:

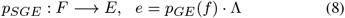

The decoder and autoencoder functions, incorporating the SVD layer, are redefined as:

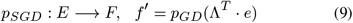

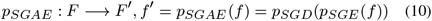

This SVD layer aids in selecting the optimal number of dimensions for CFPS, enhancing the model’s capacity for face reconstruction.

#### 5) Spiral-based Architecture

We used geometric deep learning to learn directly from the 3D facial meshes and efficiently leverage the underlying geometry by using spiral convolution operators [22]. The architecture of our geometric autoencoder is illustrated in Fig. 2. The spiral convolutional (Sconv) layer in this figure consists of first, convolving spirals on vertices of the mesh in the current layer, and second, down or up-sampling the current mesh to obtain input for the next layer. Each Sconv layer is followed by an exponential linear unit (ELU). A spiral convolution is a filter consisting of learned weights w, which is applied to a sequence of neighborhood vertices. That means,

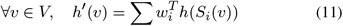

where *h*(*v*) is the input representation of vertex *v, h′*(*v*) the output representation, and *S*_*i*_(*v*) the *i*-th neighbor of *v* in the spiral [23]. The sequence was defined as a spiral around a central vertex, starting in an arbitrary direction and then proceeding in a counterclockwise direction until a fixed length was reached. In a geometric encoder based on spiral convolutions, aside from the convolution operator, a pooling operator for meshes must be incorporated. Established mesh decimation techniques used in many geometric deep learning methods reduce the number of vertices such that a good approximation of the original shape remains, but they result in irregularly sampled meshes at different steps of resolution. In contrast, we used a 3D mesh down and up-sampling scheme that retains the property of equidistant mesh sampling as defined in [24]. Starting from five initial points, the refinement is done with loop subdivision by splitting each triangular face of the mesh into four smaller triangles by connecting the midpoints of the edges. The last up-sampled mesh has 8,321 vertices and an average resolution of 2mm, meaning that the average edge length is 2mm. For our geometric encoder, the five highest levels of resolution (shown in Fig. 2) are kept, and their output is passed through the fully connected layers of our encoder. In-house experiments showed that other sampling schemes are equally effective and can be used instead. The number of spirals in each layer was chosen empirically based on the previous and related works [24], as well as in other in-house projects where similar facial data structures are used. The length of the spiral filters was set to 19 for the first two layers with the highest resolution, and a length of 6 was chosen for the following layers with lower resolution. These choices were made such that for higher resolution meshes two-ring neighbors (=19 vertices) and for lower resolution meshes one-ring neighbors (=6 vertices) are covered by a spiral filter. Larger spiral lengths were initially tested for the first layers in a geometric autoencoder and no significant improvement in reconstruction performance was observed. A shorter spiral length, covering one-ring neighbors (9 vertices), was also tested for the first layers, and the difference in performance was not significant. Therefore, to decrease the computation cost, one can choose a spiral length of nine over 16. Since we have a fixed topology enforced on all faces, the spirals were determined only once, on the template mesh.

**Fig. 2:**
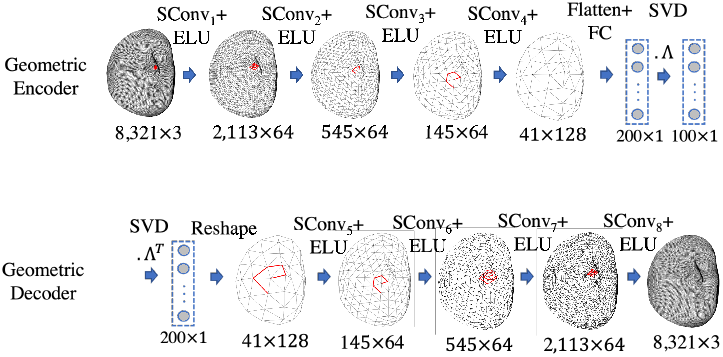
The architecture of a singular geometric autoencoder (SGAE) with a singular value decomposition (SVD) layer. Λ contains right singular vectors of the SVD. Once trained, the geometric encoder constitutes the projection function *p*_*SGE*_, and the geometric decoder constitutes the decoding function *p*_*SGD*_.

#### 6) Classification

In this study, we performed a *one vs all* classification of syndromes, where a specific syndrome is preselected, and patients are classified as either having or not having that condition, constituting a binary classification or a syndrome identification task [4], [25], [26]. We also conducted a syndrome identification task, answering the question: Given a patient, which syndrome class is most likely? To do this, a *multiclass classification* was implemented using a balanced K-nearest neighbor (KNN) classifier with *K* = 10. This classifier was employed on the projection of individual profiles into the CFPSs, derived from the DL models (TB-SGAE, TB-GAE, and TB-SGE) and baseline (as per Equations 10 and 1).)

The standard KNN algorithm calculates distances between a test image and all samples in the training set, selecting the K nearest neighbors to determine the most common label. A balanced KNN variant assigns varying weights to each training sample. This weighting counters the bias towards overrepresented classes in the dataset, similar to what was proposed by Tan et al. [27]. Specifically, the weight assigned to a sample is inversely related to the frequency of its class in the dataset, ensuring a more equitable representation of underrepresented classes in the prediction process:

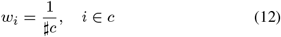

where *w*_*i*_ is the weight attributed to sample *i, c* is a group of samples from the same class (with *i* ∈ *c*) and *♯c* is the cardinality of *c*. We set K=10 since it is the minimum group size in our dataset. While one vs all classification requires the usage of a different binary classifier for each class, multiclass classification can be performed with a single KNN classifier.

Utilizing our multiclass classification results, we extract the best-S predictions to generate top-S cumulative accuracy curves. This method refines the potential diagnoses spectrum, offering more precise outcomes compared to single-label classification. Additionally, it deepens the involvement of clinicians in the diagnostic process, thus enhancing their trust in these advanced classification tools.

#### 7) Training

All models were trained on an NVIDIA GeForce RTX 2080 Ti, 64 GB RAM, with PyTorch 1.1.0. The Adam optimizer was used for 600 epochs with a batch size of 30 (limited by the maximum GPU memory), an initial learning rate of 1e-3 chosen based on experiments ran for a range of (1e-1,1e-8), and a decay rate of 0.99 was applied after each epoch.

### D. Experiments

The development of a Clinical Face Phenotype Space (CFPS) is based on the assumption that it provides a clinically meaningful and useful model for variation within and among classes. To validate this, we conducted four experiments, each examining different aspects of the CFPSs. The experiments assessed the complete network TB-SGAE against a baseline (PCA+LDA) and two network versions with specific components removed: TB-SGE (without the decoder) and TB-GAE (without the SVD layer).

**Experiment one** focused on evaluating and comparing the classification capacities of our CFPSs against a linear baseline using a 5-fold cross-validation method. In each fold, 20% of data per group was designated for testing, and the remaining 80% for training. We assessed classification performance using the top-S cumulative curves for syndrome identification using the multiclass classification, as well as sensitivity, specificity, and balanced accuracy of the one-vs-all classifier. Sensitivity measures the classifier’s ability to identify true positives, specificity assesses the identification of true negatives, and balanced accuracy averages the two. A paired two-tailed Wilcoxon signed rank test determined the statistical significance of these metrics, with distance scales of embeddings normalized for comparison. We also explored how well categories A, B, and C correlated with the classification performance of our CFPS, aiming to determine if the space accurately reflects clinical knowledge of the syndrome groups. To also visually assess these, a 2D visualization of the CFPS was generated using the Uniform Manifold Approximation and Projection (UMAP) algorithm to provide visual feedback on the CFPS structure. Furthermore, we expected that, in general, phenotypic characteristics being syndrome uniqueness (the median of distances between the average shape of a syndrome to all other average syndrome shapes), cohesion (the median shape distance, measured on landmarks, of all individuals in each group to the average shape of the group), and severity (the average shape distance between the subjects with a syndrome and the mean shape for controls [17]) should predict accuracy to a substantial degree, and this may also be impacted by sample size. Given the correlations among the phenotypic predictors, it is difficult to investigate their effects independently. To do so, predictors were combined into a single latent variable using a PLS regression of accuracy (in each space) onto the phenotypic predictors and sample size, with one latent component.

**Experiment two** investigated the ability to reconstruct faces from the CFPS using training and out-of-fold (OOF) error measurements. The training error, or reconstruction loss, gauged the model’s efficacy in capturing shape variation. The OOF error, calculated as the mean absolute error for test set samples, reflected the model’s performance with unseen data.

In **Experiment three**, we assessed the generalization capability in clustering syndrome groups not included in CFPS training. Six syndrome groups with distinct phenotypic features were omitted during training. Their post-training projection into the space was analyzed through the determination of the clustering improvement factor (CIF). CIF, which measures the clustering improvement over random distributions, was computed using the expected and observed ranks of the nearest positive matches. Comparisons were made between the complete model (TB-SGAE), the baseline (PCA+LDA), and an unsupervised linear approach (PCA).

**Experiment four** verified known relationships between specific syndrome groups within the CFPS. Here, we focused on Noonan, Costello, Cardiofaciocutaneous syndrome, and Neurofibromatosis Type I (NF1), known collectively as RASopathies. A p-value for the average distance between cluster centers of these groups was calculated against an empirical null distribution generated by randomly selecting four groups from the dataset 10000 times. We also show the positioning of RASopathies within the 2D UMAP visualization of the CFPS.

Through these experiments, we aim to rigorously evaluate the CFPS’s performance in various aspects, including classification performance, facial reconstruction, generalization to unseen data, and preserving the relatedness between syndromes, thus providing insights into the developed CFPS’s effectiveness and clinical utility.

## III. Results

For the first experiment, Figure 3a demonstrates the syndrome identification capacity of the CFPS, in top-S accuracy curves obtained from multiclass classification. These metrics are derived in an individual-based manner, meaning that the average values are calculated across all the test samples. This figure is generated for *S*≤10 in TB-SGAE, TB-GAE, TB-SGE, and the baseline. The DL methods outperformed the baseline across all *S* ≤ 10 values, with TB-SGAE showing the highest performance. A notable performance gap was observed at lower *S* values, with a convergence trend in accuracy as *S* increases. Figure3b offers a detailed analysis for TB-SGAE, highlighting its effectiveness in identifying controls and varying accuracy across different syndrome categories, following clinical expectation, with convergence in accuracy for higher *S* values.

**Fig. 3:**
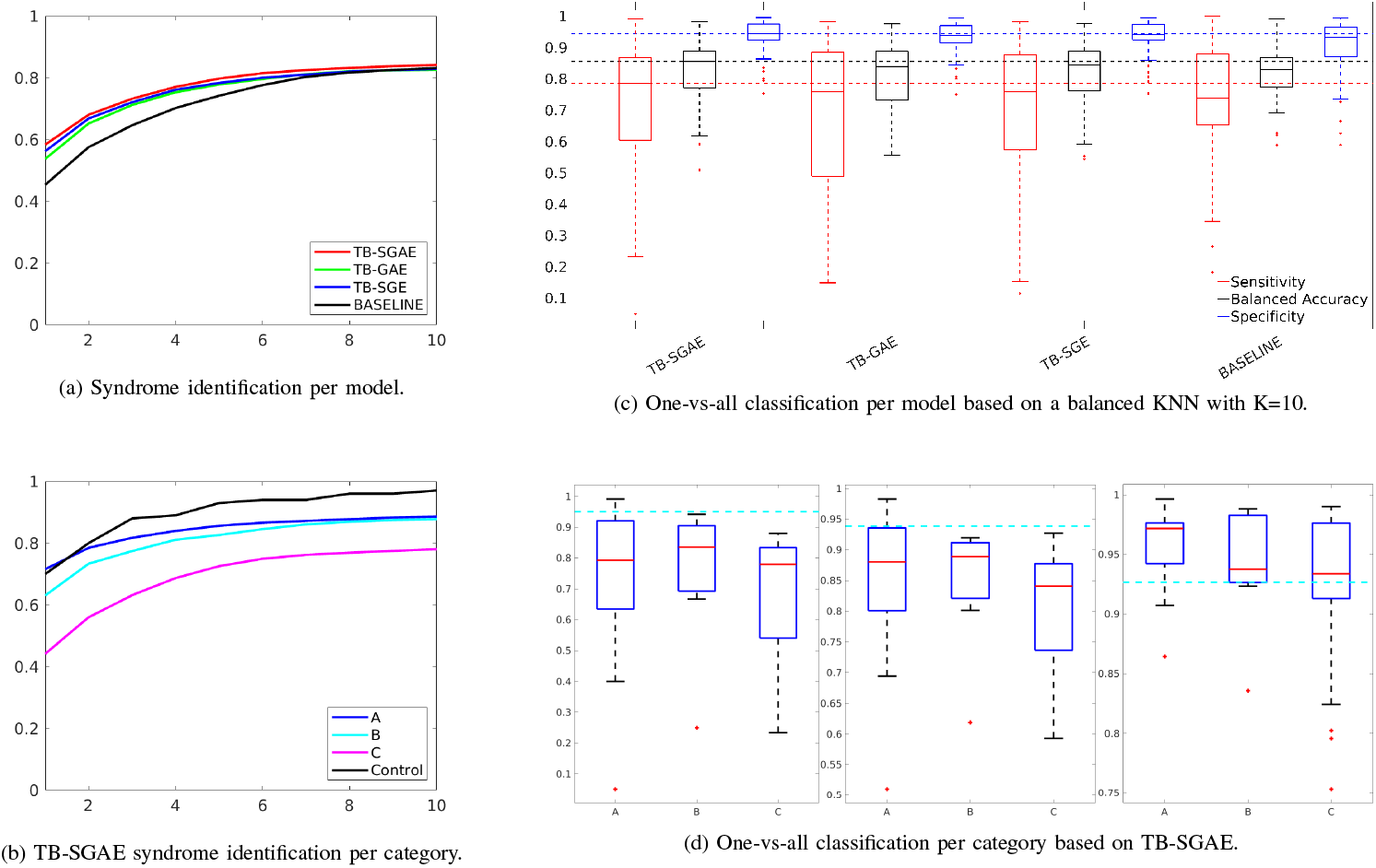
Experiment one - Classification Performances: (a) Individual-based top-*S* accuracy plots for TB-SGAE, TB-GAE, TB-SGE, and the baseline (*S* ≤ 10); (b) Individual-based top-*S* accuracy plot for TB-SGAE (*S* ≤ 10) separated by categories (A, B, C, and controls); (c) Classification metrics based on a balanced KNN classifier with *K* = 10; (d) The average group-level metrics for all groups in categories A, B, and C based on TB-SGAE.

In addition to the syndrome identification, Table 2 reports the average and the standard error of the one-vs-all classification measures over the five cross-validation folds. These results are based on the CFPS obtained by TB-SGAE, TB-GAE (SVD layer removed), TB-SGE (decoder removed), and the baseline (PCA+LDA). These metrics are again derived in an individual-based manner. In contrast, the distributions of metrics, averaged within each syndrome group over the five folds are shown using boxplots in Fig. 3c. These results are determined in a group-wise manner, meaning that they are based on the average metrics’ values for each syndrome class. The p-values of the statistical test comparing TB-SGAE and the baseline were 0.6078, 0.0499 and 4.02e-7 for sensitivity, balanced accuracy and specificity, respectively. According to the results, the performance of the TB-SGAE was significantly higher than the baseline (p-value*<*0.05) for all metrics, except for the sensitivity, where the statistical difference was not significant. Fig. 3c and Table 2 also compare TB-SGAE with TB-SGE and TB-GAE to investigate the contribution of the decoder and the SVD layer, respectively. The decoder significantly improved the classification measures. Removing the SVD layer significantly decreased all performance indicators.

**TABLE II:** Comparison of discriminating individual-based metrics computed by applying a balanced KNN classifier to the embeddings extracted by TB-SGAE, TB-SGE, TB-GAE and the linear baseline (PCA + LDA). The reported metrics are sensitivity, balanced accuracy and specificity. The best value achieved for each metric is in bold.

| Model | Sensitivity | Balanced Accuracy | Specificity |
| --- | --- | --- | --- |
| **TB-SGAE** | **0.8234 ± 0.1920** | **0.8725 ± 0.0918** | **0.9216 ± 0.0597** |
| **TB-GAE** | 0.8101 ± 0.2043 | 0.8652 ± 0.0988 | 0.9202 ± 0.0560 |
| **TB-SGE** | 0.8110 ± 0.2095 | 0.8655 ± 0.0993 | 0.9200 ± 0.0591 |
| **BASELINE** | 0.8223 ± 0.1721 | 0.8427 ± 0.0822 | 0.8630 ± 0.1146 |

Fig. 3d shows the distributions of TB-SGAE metrics, averaged within each syndrome group over the five folds, stratified by the clinical categorization (A, B, or C). Syndromes in categories A and B had a higher median sensitivity, balanced accuracy and specificity than those in category C. While category B showed slightly higher sensitivity and balanced accuracy values, A largely surpassed it in terms of specificity, which is consistent with clinical expectations behind the categorization of the syndromes.

Fig. 4b shows the 2D visualization of the CFPS obtained from TB-SGAE using the UMAP algorithm. The projection of individuals in the train set (larger dots) and test set (smaller dots) are colored by their categorization. The PLS regression of balanced accuracy onto phenotypic predictors and sample size is shown in Fig. 4a. The standardized coefficients of the linear combination and the regression of accuracy onto the derived latent variable are also shown. Accuracy in the CFPS of the TB-SGAE and TB-SGE is significantly predicted by the phenotypic measures and sample size, when compared to the baseline (PCA+LDA). This observation also underscores the significance of sample size effect in deep learning models compared to the linear method, encouraging the collection of additional data in future studies. In addition, it is noteworthy that cohesion appears to be a less robust predictor in TB-SGE compared to its predictive relevance in TB-SGAE. This suggests a potential relationship between the model’s face reconstruction capability and the utilization of cohesion as a predictive factor for facial shape variations within distinct groups.

**Fig. 4:**
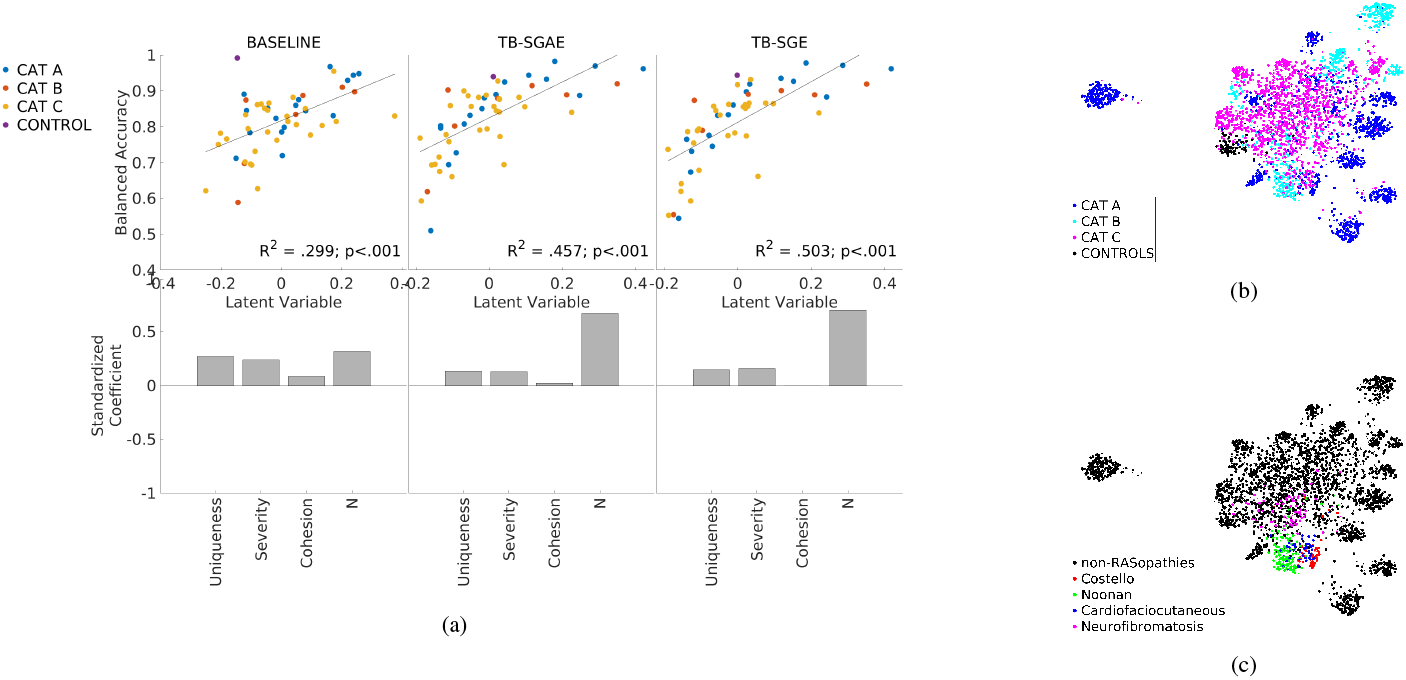
(a) PLS regression of accuracy onto phenotypic predictors and sample size. (b) 2D UMAP visualization of the trainset (smaller dots, and test set (larger dots) into the space, colored by categories. (c) Colored 2D UMAP visualization of the four RASopathies together with the rest of the trainset (smaller dots) and test set (larger dots) colored in black.

In experiment two, the training and OOF error of reconstruction for TB-SGAE were 0.1597 and 0.1705, respectively. Reconstruction error per vertex is shown in Fig. 5a. The error bar is scaled in millimeters. The average error per vertex was less than 1 mm. Nevertheless, the heatmaps indicate that regions around the mouth, nose, and eyes had relatively higher errors. The lips and mouth regions are sensitive to expression variation, introducing extra complexity for the model to learn. To visually assess the precision and smoothness of the reconstructions from the CFPS of the complete model, the average test-set projections can be reconstructed. Fig. 5b shows reconstructions for four groups of Achondroplasia, Wolf Hirschhorn, Apert and Williams from Category A syndromes.

**Fig. 5:**
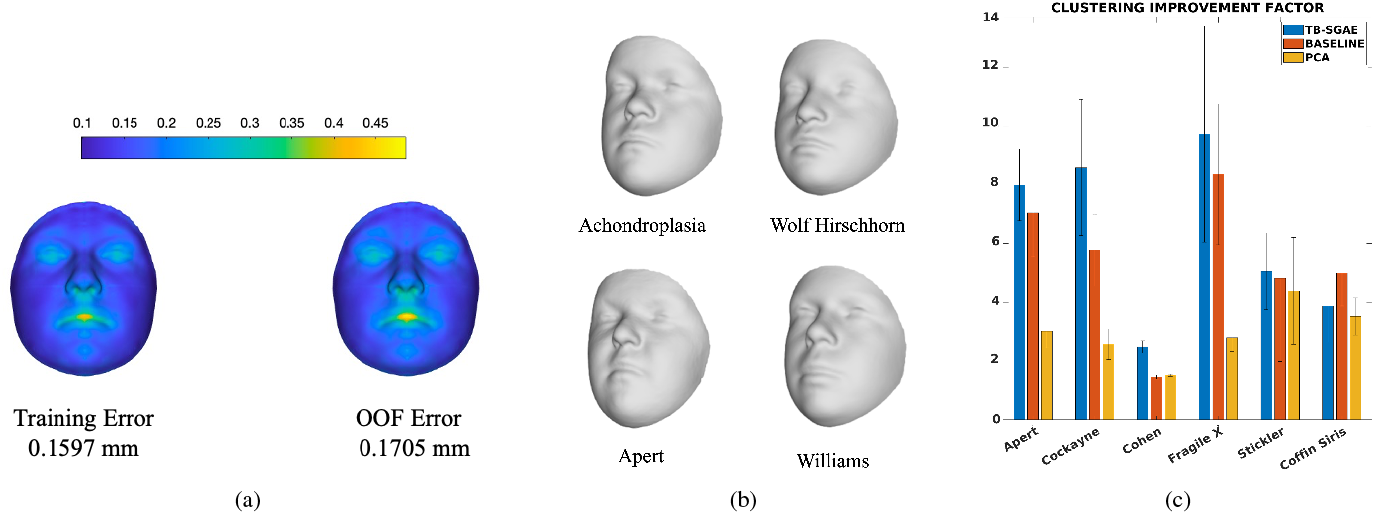
(a) The training error of reconstruction (left) and out-of-fold error of reconstruction (right); (b) The reconstruction of the average embedding of individuals with Achondroplasia, Wolf Hirschhorn, Apert, and Williams using the geometric decoder; (c) The comparison of the clustering improvement factor (CIF) for individuals in the six left-out groups of the generalization test (experiment 3), projected to the CFPSs obtained by TB-SGAE, PCA+LDA (baseline), and PCA. Error bars indicate the standard error of the mean over five folds.

The third experiment investigated the ability of the space to generalize to unseen syndromes. Six groups of syndromes were left out from the training set, and the CFPS was trained based on the 47 remaining groups. The unseen groups and the test set were merged and projected to the space. Based on these projections the CIF were computed. Fig. 5c shows the average results for TB-SGAE, the baseline (PCA+LDA) and PCA over five folds of data. The CIF was higher for five out of the six syndromes for TB-SGAE. The (unsupervised) space of principal components had a generalization power higher than a random performance, which indicates that unsupervised facial structure or similarity already improves the clustering of unseen syndromes. In addition, applying the LDA transformation to the PCA scores increased the CIF of all the syndrome groups but Cohen syndrome, which indicates that supervision further improves the clustering of unseen syndromes. Note that the maximum dimensionality of the LDA-based CFPS was bounded by the maximum number of classes in the training set minus one (i.e. 46 in this study). Therefore, we further investigated and compared the CIF results using a TB-SGAE space with 100 dimensions (used before and determined based on an SAE, see above) and 46 dimensions (equivalent to the LDA-based baseline), and observed no statistical difference (p=0.84). This is not entirely unexpected thanks to the SVD layer, which results in the most variance being coded in the lower components, making it easier to reduce dimensionality, with the minimum loss of data information.

For the last experiment, the UMAP plot in Fig. 4c shows the RASopathies grouped at the center of the UMAP, confirming these groups’ proximity in the CFPS. The statistical test also indicated that within the CFPSs based on TB-SGAE and the baseline (PCA+LDA), the average distance between testset RASopathies cluster centers in the normalized CFPSs was lower than 78.39% and 65.31% (respectively) of random selections of four groups of testset samples.

## IV. Discussion

In this work, we build a CFPS that models the range of facial dysmorphism present in 52 syndromes alongside general facial variations from a group of controls. To this end, we proposed a triplet-based singular geometric autoencoder for multi-task learning, to simultaneously learn facial shape variation and reconstruction, in an unsupervised way, and group discriminations with the supervision of syndrome labels.

The existing CNNs for syndrome classification or building CFPSs are mostly based on large-scale 2D photographs of patients with genetic syndromes. By now, large-scale databases of 3D photographs of clinical populations have been collected. Considering the expected growth in the popularity and accessibility of portable 3D imaging hardware, building systems that apply to this imaging modality is essential to fully exploit the 3D shape information contained in such images. With the recently developed field of GDL, CNNs are now directly applicable to 3D images. This eliminates the need for any domain transformation. Therefore, in this work, we aimed to build a CFPS based on 3D facial images using spiral convolutional operators with which we facilitate both syndrome classification and facial reconstruction. Once learned, we evaluated the main properties of the CFPS, such as the classification of syndromes, generalization to novel syndromes, and the recapitulation of related genetic diseases. We also assessed the reconstruction precision from the CFPS and investigated the phenotypic shape predictors of the classification. We compared the performance of our space to a linear baseline which consists of PCA for dimensionality reduction and LDA, a linear metric learner. Similar work on 2D data [6] estimates the factor by which clustering is improved compared to random chance (CIF). Compared to such random performance, LDA is a much more difficult baseline to match or improve on. In fact, for statistical shape analysis, LDA and its regularized variants were and still are strong and popular methods that are also used and outperformed many other classifiers in the 3D syndrome classification published in [17].

Our proposed model consists of three main components. The first is a triplet-based encoder which was used in the recent syndrome classification work in [26] to optimize the distances among individuals belonging to different syndrome groups. In the triplet-loss function, the focus is on learning the CFPS such that the distances are a measure of similarity and group membership and therefore it contributes to the classification power of the space. The second component is a decoder that not only allows the reconstruction of a face from an embedding in a CFPS but also improves the classification performance of the system. The third component, being an SVD layer, makes it simpler to select the dimensionality of the space without retraining and also improves the classification, reconstruction, and generalization aspects of the CFPS.

We showed that the CFPS built based on the complete model (TB-SGAE) outperforms the classification performances of the linear model which consists of PCA for dimensionality reduction and LDA for metric learning. More specifically, TB-SGAE presented the highest classification performance in all the considered metrics (being sensitivity, balanced accuracy and specificity in one-vs-all classification and the syndrome identification task assessed through top-*S* accuracy in multiclass classification) when compared to all the tested models. This illustrates the usefulness of both the SVD layer (by comparison with TB-GAE) and the decoder (by comparison with TB-SGE). Although a direct comparison with the state-of-the-art 3D syndrome diagnosis in [17] is not available, the classification results are competitive.

Furthermore, we investigated whether the CFPS is inline with the expectations of clinicians. Experts in clinical genetics categorized syndromes into three classes. Two include syndromes that are phenotypically distinctive (A and B) and the third class of syndromes that are not necessarily phenotypically distinctive (C). Graphically, the UMAP visualization (Fig. 4b) shows that syndromes in categories A and B generate more isolated and clear clusters around the corners while category C groups show less clear cluster boundaries and are positioned close together around the center of the mapped embeddings. Category A and B are also more distant from controls than category C. This observation is in line with the high performance of Category A and B over category C in the syndrome identification test (Fig. 3b), as well as the one-vs-all classification performance (Fig. 3d).

With the ability to construct a face from an embedding, we can visualize and hence explain and understand the embeddings better. The decoder component facilitates the reconstruction of encoded facial shapes with less than 1 mm error. In addition, thanks to the orthonormal dimensions, we have a coordinate system in the space that properly spans every vector and thus allows us to explore and interpolate the space more structurally. In other words, one can manipulate one dimension without changing the values on other dimensions. This property of a vector basis or coordinate system is not available in a default autoencoder.

Furthermore, we evaluated the clustering generalization onto novel syndromes that have not been included in the training set. We computed and compared the CIF for the novel syndromes. This comparison shows that the clustering improves from a random chance for both baseline and TB-SGAE. Compared to the linear baseline, the improvement is stronger for five out of six novel syndromes within our CFPS. The comparison of the generalization power between the supervised linear metric learning approach (PCA+LDA) and that of the unsupervised and linear PCA shows that in five out of six left-out groups supervised learning improves the performance. The group that has superior performance with PCA (Cohen syndrome) is known to be clinically difficult to recognize from the face, and this observation suggests that the supervision of metric learning has less influence on groups that have little to no facial clues for diagnosis. It is also worth mentioning that the CIF obtained by PCA suggests that despite the unsupervised nature of this method, it still is powerful enough to improve the clustering factor considerably compared to random performance.

Finally, we tested for expected low distance among the four RASopathies, an etiologically related group of disorders caused by mutations in genes encoding the RAS/MAPK pathway. Results of the statistical test indicated that the four groups are close together in the CFPS of TB-SGAE. This test provided us with one piece of evidence towards the recapitulation of the relatedness in the CFPS, although it does not demonstrate any improvement of the deep metric learners over the linear baseline, in which our measure of this recapitulation was already close to the ceiling. When genetic data are available, more comprehensive tests, correlating measures of genetic similarity to phenotypic distance should be performed. For example, genetic similarity can be based on protein-to-protein interaction as per Ferry et al. [6] or distance based on patterns of DNA methylation [28].

## V. Conclusion

In this work, we proposed a CFPS learner based on 3D facial images and GDL techniques for large-scale syndrome analysis. The proposed model consists of the base component being a geometric encoder, which is further expanded by our additional components being a geometric decoder, with which high-precision facial shapes are reconstructed from an embedding in the CFPS, and a singular value decomposition layer to encode a structured facial mesh into an orthonormal 100-dimensional CFPS. We used a multi-task learning approach to train the model in an end-to-end manner. The loss function combines the supervised triplet-loss function with the unsupervised reconstruction-loss. In summary, we showed that supervised and unsupervised learning strategies both improve the clustering factor compared to a random performance. Moreover, supervised learning leads to superior performance compared to unsupervised learning only. Lastly, the proposed GDL-based model learns a CFPS that outperforms the linear metric learning baseline (consisting of PCA and LDA), in both syndrome classification and generalization to novel syndromes. We proved the contribution of each added component in the classification and reconstruction capacity of the CFPS. More precisely, we showed that the attached decoder not only facilitated the ability to reconstruct patient faces and generate synthetic faces but also improved the classification performance of the model both in one vs all and multiclass frameworks. In addition, the orthonormal base of the CFPS facilitated by the SVD layer has considerably impacted the classification performance, both in one vs all and multiclass frameworks. We also showed that the space strongly replicates clinical expectations such that the classification measures obtained from the CFPS relate to the categorization of syndromes. Furthermore, the proximity of the four RASopathies, characterized by mutations in functionally related genes, is reflected in the CFPS.

The resulting CFPS can potentially narrow the search space for diagnosing new instances of the syndromes that it represents, objectively assess facial similarity between undiagnosed patients who share a rare and novel disorder, and facilitate targeted sequencing of genomic regions to identify causal variants.

## Footnotes

1 https://www.facebase.org/

